# How do neuroglial cells respond to ultrasound induced cavitation?

**DOI:** 10.1101/2020.10.14.339002

**Authors:** Alex H. Wrede, Jie Luo, Reza Montazami, Anumantha Kanthasamy, Nicole N. Hashemi

**Affiliations:** Department of Mechanical Engineering, Iowa State University, Ames, Iowa 50011, USA; Department of Biomedical Sciences, Iowa State University, Ames, IA 50011, USA

**Keywords:** cavitation, microbubbles, ultrasound, microfluidics, traumatic brain injury

## Abstract

Reactive astrocytes are known to play a vital role in the overall response of the brain during a traumatic brain injury (TBI). Modern studies have speculated the existence of cavitation in the skull during a TBI, which has alarming potential to cause detrimental damages. Previous studies have confirmed the upregulation of various harmful genes in neurodegenerative diseases. Studying the longitudinal presence of these harmful genes in response to cavitation allows for optimized understanding and treatment methods in cavitation exposure. We seek to characterize the longitudinal genetic expression levels that astrocytes exhibit after exposure to cavitation and further elucidate the startling presence of cranial cavitation. Astrocytic expression levels of various common genes that have been documented in TBI research are our target of interest, like, TNFα, IL-1β, and NOS1. Results summarize specific gene trends from 1-48 hours after cavitation. Our data concludes that maximum expression is not consistently exhibited immediately after cavitation exposure, and most genes have individualized genetic trends. IL-1β shows a decreasing expression over 48 hours, and TNFα shows upregulation until the 6 hour time point but then begins to decrease in expression. The upregulation of NOS1 has been documented in neurodegenerative diseases, like Alzheimer’s and Parkinson’s disease. This study has shown a consistent upregulation in NOS1 expression from 0-48 hours. These results postulate a possible linkage between cavitation damage and neurodegenerative diseases. This analysis also provides novelty in optimizing treatments for astrocytic function post-TBI and legitimizing the concern of cranial cavitation existence. These results add motivation for future studies of cavitation elimination or minimization via advanced helmet and airbag engineering.

## 1. Introduction

Despite an abundance of recent efforts in TBI research, there are still undiscovered attributes in the neuronal response to these situations. Reactive astrocytes play a vital role in a variety of neurodegenerative diseases, including Alzheimer’s disease and Parkinson’s disease[1–3]. Previous studies have shown that the effects from altered astrocytic gene expression is dependent on the specific inducer of the trauma, adding an increased level of complexity in TBI response characterization. For example, ischemia induced astrocytic change represents a genetic phenotype that is beneficial and sustaining to surrounding neuronal anatomy. On the contrary, an inflammatory exposure, like lipopolysaccharide (LPS), induced astrocytes that cause detrimental gene alterations [4].

For simplicity, scientists have categorized detrimental reactive astrocytes as ‘A1’ astrocytes and protective reactive astrocytes as ‘A2’ astrocytes[3]. Ischemia and LPS are two possible exposures that will induce reactive astrocytes, but it remains a question as to how reactive astrocytes respond differently to other types of TBI exposures. A modern theory postulates that a likely inducer of reactive astrocytes following a blast-TBI is the presence of cavitation[5].

Cavitation is a byproduct of low pressure regions in the brain, which are a result from an initial compressive wave reflecting off the opposite side of the skull[6, 7]. The localized force from cavitation leaves behind profound damages in other applications, like boat propeller and hydrodynamic pump erosion[8, 9]. Understanding whether surrounding cavitation exposure produces A1s, A2s, or a combination of each, is critical in order to further unmask the complexity of TBIs. Previous research has analyzed the specific astrocytic gene expression alterations immediately after experiencing surrounding cavitation. These results showed an upregulation in a number of classic A1 specific genes, including TNFα, IL-1β, C1q, Serping1, NOS1, and IL-6. Results showed minimal changes in classic A2 specific genes with the exception of CD14[10].

In this study, we expand on previous methods to conduct a longitudinal study that specifically elucidates astrocytic gene expression abnormalities over time, after experiencing exposure to surrounding cavitation. It is known that gene expressions can change over time and further characterizing this trend in reactive astrocytes has great potential[4], like leading to advanced pharmaceutical techniques to restore altered cellular function. This study analyzes the astrocytic gene expressions up to 48 hours post-cavitation exposure via quantitative polyermase chain reaction methods (qPCR) methods. These genetic analytics yield insight on the behaviors that reactive astrocytes exhibit after exposure to nearby cavitation and provide a foundation of knowledge that helps further characterize reactive astrocytic nature in TBIs.

## 2. Materials and Methods

### 2.1. Cell Model

Mouse astrocytes (CRL-2541, ATCC, Manassas, VA) are solely used in this project because of their documented utility in TBI studies[11, 12]. This particular cell line is a clonal permanent cell line in which the cells have astrocytic properties. The astrocytes were cultured according to company protocol in base media (ATCC, 30-2002). The growth media was also composed of 10% FBS and 1% penicillin (10,000 U/mL)-streptomycin (10,000 μg/mL) (Gibco, Waltham, MA). All cells in this study were from culture passage numbers 5-7. The astrocytes were stored in an incubator set with constant parameters of 37 °C and 5% CO_2_. The mouse astrocytes were seeded into the 6-plates at a concentration of 9.04 * 10^6^ cells per well and stored for 24 hours in the incubator prior to experimentation.

### 2.2. Cavitation Apparatus

The method of creating controlled cavitation is similar to that of previous studies[13–15]. The main difference in the apparatus configuration of this experiment is the cells are seeded onto sterilized 6-well culture plates instead of using microfiber scaffolds, shown in Figure 1. All components are housed in a 5.7 liter tank filled with phosphate buffered saline (PBS). During trials, the experimental samples are directed downward toward capillary tubing (Polymicro Technologies, 1068150003) that is continuously releasing microbubbles (MBs) ~60μm in diameter at a rate of ~150 MBs/min. The 6-well plate is mounted on a 3-axis adjustable stage (MT1, Thor Labs, Newton, New Jersey) allowing for MBs to adhere arbitrarily across the cell-seeded surface. After a 350-400 MBs are adhered to each well the culture plate is then rotated 180°, exposing the astrocyte-MB complex to a 100 kHz transducer positioned overhead in the tank. The transducer induces resonant conditions causing the MBs to oscillate and fragment, also known as induced cavitation (center frequency 100 kHz, 5 cycles, pulse repetition frequency 59 Hz, 260 Vpp). The frequency signal is amplified using a broadband power amplifier (E&I, 1140LA). The control samples were introduced to an interchangeable environment, but they did not experience nearby cavitation because there were not any MBs arbitrarily captured in these wells.

**Figure 1:**
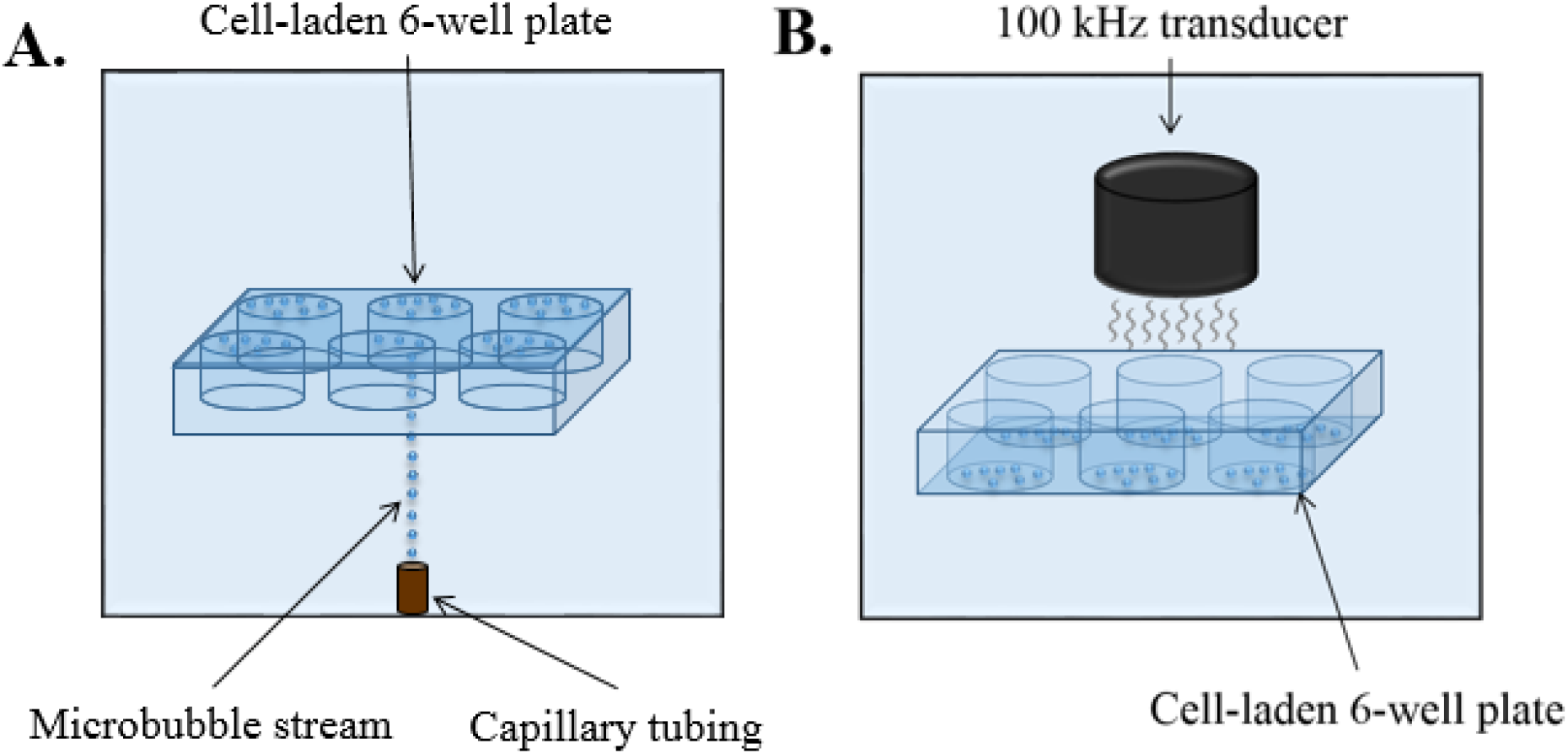
Apparatus layout of MB capturing and induced cavitation on cell-laden 6-well plates. (A) 350-400 MBs are captured and adhered to the bottom of all the cell-laden culture wells. (B) The 6-well plate is rotated 180° and exposed to the 100 kHz transducer. The transducer is centered above each well individually. When the transducer is turned on the astrocytes are introduced to surrounding cavitation.

### 2.3. qPCR Analysis

A full 6-well plate was designated for a specific time point, three wells for experimental samples and 3 wells for control samples. Designating one plate for each time point was advantageous because the experimental and control samples experienced exact amounts of PBS and ultrasound exposure with this orientation. Another advantage of this setup is the ability to retain cells that detached from the well due to the cavitation force. When the cells detach they drift within the boundaries of each specific well, allowing them to be part of the post-cavitation analysis. Previous methods use cell-laden coverslips without ridges so the detached cells are unable to be collected[10]. After the plate was pulled from the tank, the remaining PBS solution was pipetted out and replaced with fresh growth media. The pipetted PBS was centrifuged at 1500 rpm for five minutes to collect all the detached cells. The pelleted cells were put back in the original wells and placed in the incubator until its specific designated time point after exposure. Cell response was studied 1 hour, 3 hours, 6 hours, 12 hours, 24 hours, and 48 hours after cavitation exposure.

The cell-laden 6-well plates were trypsinized and combined to create one control and one experimental sample at each designated time point. Each sample was then centrifuged at 1500 rpm for 5 minutes to pellet the cells. The cell pellet was broken with TRIzol reagent (Invitrogen Cat number: 15596026) and frozen at −80 °C until all time points were resuspended. Homogenization was done in TRIzol RNA isolation and specifically conducted following the TRIzol reagent protocol from the manufacturer. The entirety of the qPCR process in obtaining relative gene expression values is consistent with that of previous studies[10, 16].

## 3. Results and Discussion

Longitudinal gene expression analysis has been done on both A1 and A2 genes from 1-48 hours in mouse astrocyte cells. Figure 2 highlights the expression trends for A1 genes, including, Serping1, C1q, TNFα, NOS1, IL-6, and IL-1β. All data is represented as a fold change that is normalized to a control sample that was absent of cavitation damage. This data collection is an exploratory approach to elucidate any noticeable trends in expression values. Combining these trends with previous data that quantitates the expressions immediately after cavitation allows for a full 48 hour timeline[10]. This longitudinal response is novel for many reasons, including elucidating further insight on the overall biological response of the brain following cavitation exposure.

**Figure 2:**
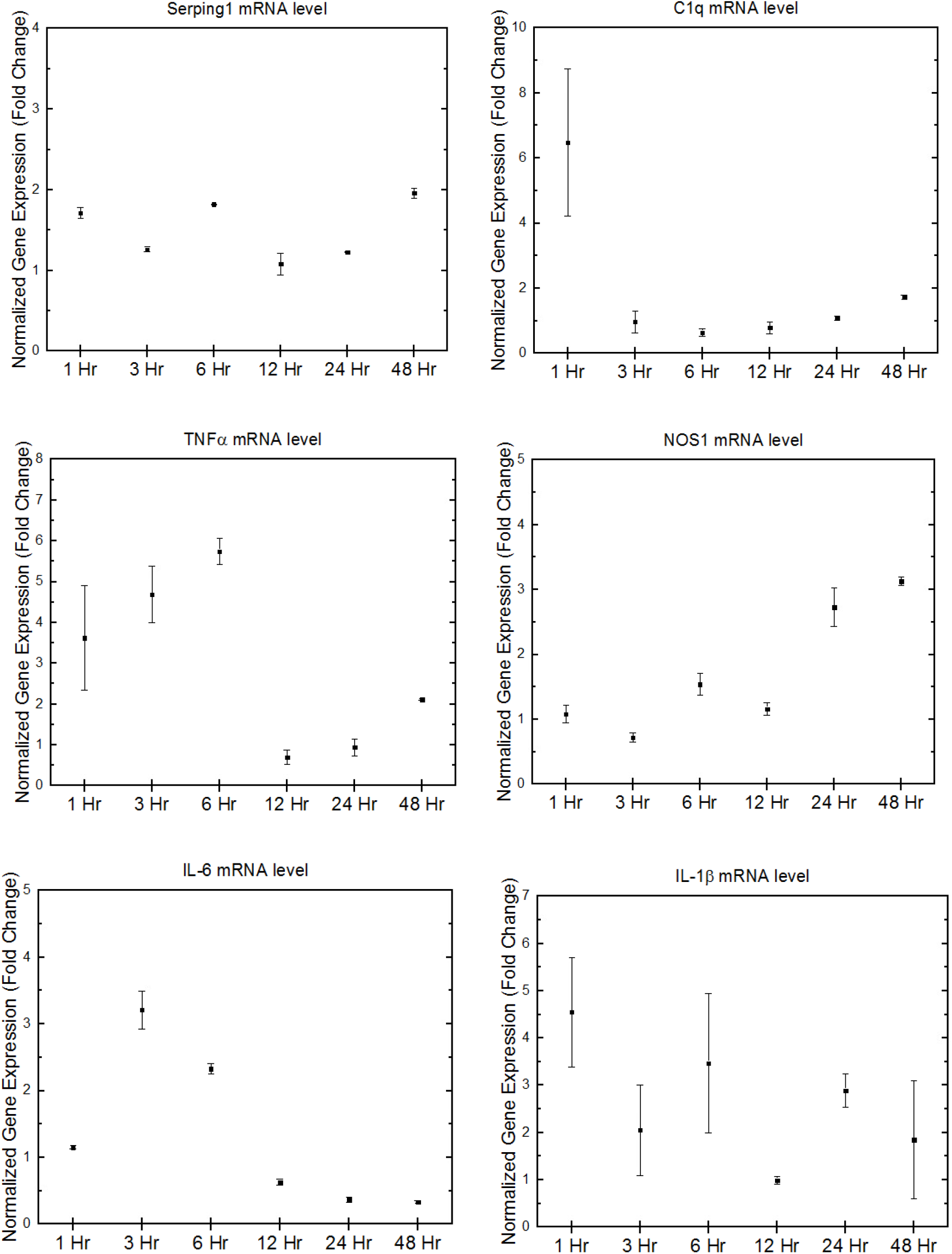
Longitudinal gene expression summaries of A1 genes. Error bars represent standard error of the mean. N = 9.

Previous data found Serping1, IL-1β, and C1q having a +5 fold change immediately after cavitation[10]. Figure 2 illustrates Serping1, IL-1β, and C1q all showing upregulation one hour after cavitation decreasing with time over 48 hours. Serping1 decreases rapidly and is shown to consistently remain within a 1-2 fold change from 1-48 hours. C1q shows similar trends with the exception of a significant upregulation remaining at 1 hour. IL-1β shows a decreasing trend over 48 hours but has a larger amount of variance and upregulation. TNFα and IL-6 both show upregulation over the first time points but at the 6 hour and 3 hour time points they begin to decrease in expression, respectively. A similarity found in the A1 genetic trends is that every gene expresses levels similar to that of the control after 48 hours, with the exception of NOS1. From the A1 genes of interest, NOS1 is the only A1 gene that represents an increasing expression trend throughout 1-48 hours. A variety of neurodegenerative diseases, like Alzheimer’s and Parkinson’s disease, have been linked to upregulation of NOS1[2]. The nature and symptoms from these diseases also escalate over time, ultimately motivating a hypothesis that directly associates cavitation exposure with Alzheimer’s and Parkinson’s disease. Figure 3 summarizes the gene expression trends for some A2 genes, including, tm4sf1, sphk1, CD14, and Arginase1. The overall trends of sphk1 and Arginase1 represent similar patterns, increasing in expression until peaking at 6 hours and showing a ≤ 1 fold change thereafter. Previous data concluded CD14 having +5 fold change immediately after cavitation[10]. This study demonstrates an overall trend of CD14 decreasing with time. Tm4sf1 shows consist expressions compared to the control, with the exception of a +2 fold change at 24 hours.

**Figure 3:**
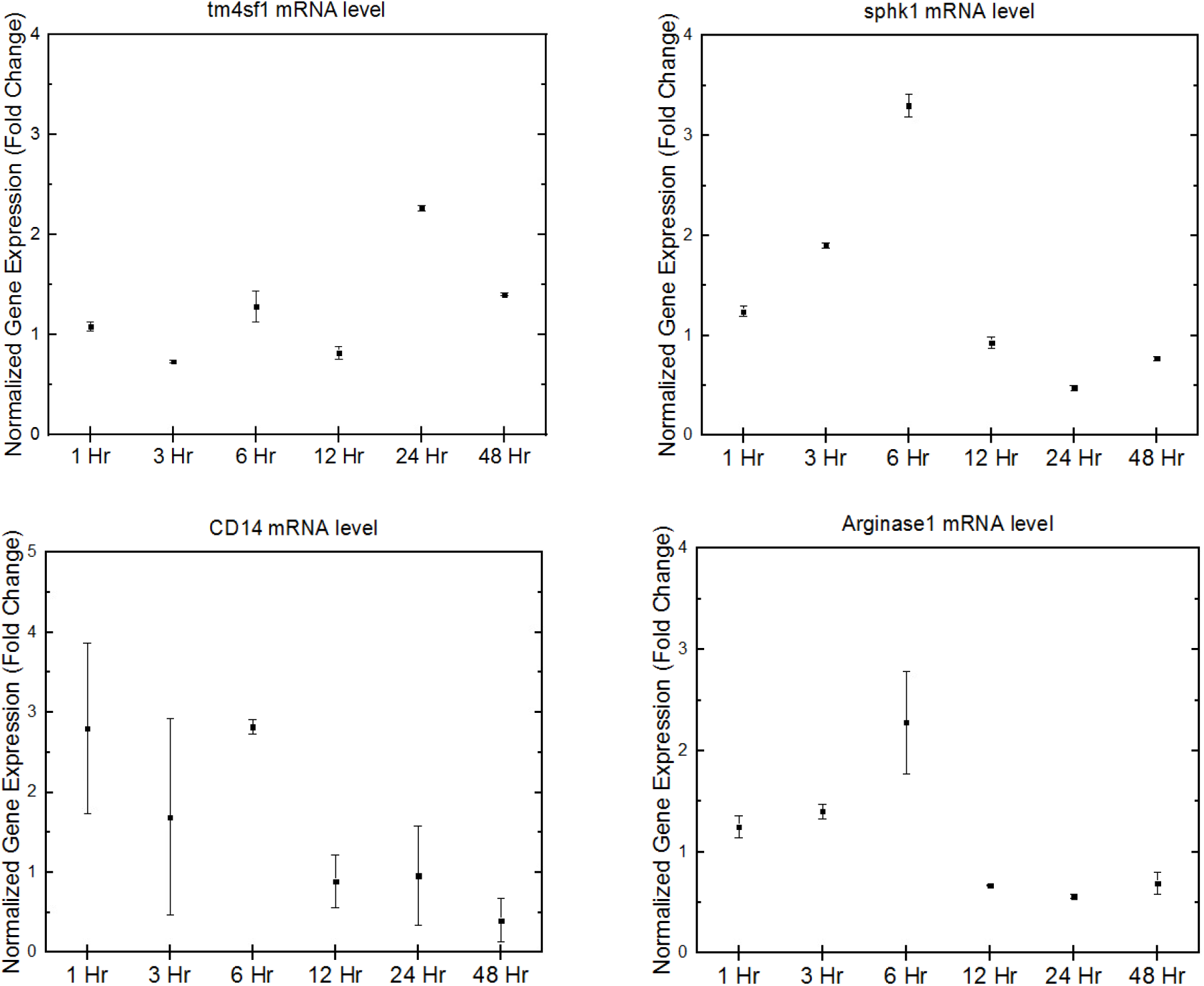
Longitudinal gene expression summaries of A2 genes. Error bars represent standard error of mean. N = 9.

This exploratory study has illustrated the unique response that cavitation induces on various A1 and A2 genes. These are the beginning trials of this investigation and future research aims to conduct RNA sequencing analytics to highlight larger groupings in the entire genome that experience expression alterations. The selected genes of this study were chosen based off of significance in existing literature. RNA sequencing will expose additional groups of genes that are of interest. Postulating a method of treatment has some value with the collected data of this study, but it is important to remember that cavitation is one of multiple injuries that occur during a TBI. Studying the genetic trends from overall TBI exposure has been heavily researched and treatment techniques have been proposed[17]. The genetic responses gathered in this study help raise awareness of the distinct changes that astrocytes undergo from solely cavitation exposure. The conditions in which cavitation likely exists in the human skull is, for the most part, unknown so we are proposing further studies that focus on characterizing the parameters that allow cavitation to exist in a TBI situation. The ultimate goal would be to take the parameter limits and engineer helmets and automotive airbags to keep the conditions within the skull from reaching these thresholds. Using these proposed parameter limits in conjunction with the longitudinal results from this study, treatments can be given at the optimized time point to assist in cranial recovery if the skull was simulated through a cavitation-inducing environment. All in all, cavitation existing in the skull urges the motivation for many future studies. The longitudinal genetic trends in this study are novel in helping piece together the complexity of the unique response that the brain has to cavitation.

## 4. Conclusions

Optimizing the specific diagnosis and care for TBIs is currently a major challenge. This is due to the fact that TBIs are a complex injury that have a variety of parameters to account for. Recent studies have added an additional layer to this puzzle with the idea that cavitation exists inside the skull during a TBI. Studying the response that neuronal anatomy has to this event is vital in optimizing TBI diagnosis and care. This study has documented the longitudinal gene expression trends that exist in a number of common A1 and A2 genes. Results have shown that each A1 and A2 gene of interest has a unique expression response and peaks at varying time points. These trends are different than a typical neurodegenerative (A1) or neuroprotective (A2) astrocyte, and hence we are postulating that cavitation induces a unique astrocytic phenotype. Treatments should be optimally given prior to the discovered peaks of the A1 genes to reduce damages that are known to be present in neurodegenerative diseases. These findings are novel in the progression of TBI knowledge and help illustrate more of the nature in the response of the brain in TBI situations. Future studies aim to expand on these results and gather complete genome trends.

## Acknowledgement

This work was supported by the Office of Naval Research (ONR) Grant N000141612246 and ONR Grant N000141712620. The authors thank Timothy Bigelow for his help with ultrasound logistics and lending a power amplifier.

The data that support the findings of this study are available from the corresponding author upon reasonable request.

